# Revival of light-evoked neuronal signals in the post-mortem mouse and human retina

**DOI:** 10.1101/2020.06.30.180497

**Authors:** Fatima Abbas, Silke Becker, Bryan W. Jones, Ludovic S. Mure, Satchidananda Panda, Anne Hanneken, Frans Vinberg

## Abstract

The retina, a highly metabolic tissue in the central nervous system, consumes the most oxygen and energy stores in the body by tissue mass/volume. Consequently, it is not surprising that retinal ischemia leads to a rapid loss of retinal light responses and electrical transmission. In this study, we show that despite a swift decline of retinal light responses after circulatory death (decay time constant τ = ∼1-2 min), we were able to restore mouse rod and cone photoreceptor light signals from enucleated eyes up to 3 h postmortem with significantly better postmortem recovery of cone versus rod light responses. We also demonstrate that both rod and cone phototransduction are more resistant to postmortem enucleation delay than synaptic transmission to second order neurons (bipolar cells). Encouraged by these analyses and the lack of previously successful postmortem retinal light recordings from human foveal/macular photoreceptors, we attempted to restore light responses in the human macula using donor eyes harvested 0.5 – 5 hours postmortem. Here we show successful recordings of human macular cone photoresponses in samples obtained from eyes enucleated up to 5 hours postmortem. Comparing ‘freshly’ enucleated non-human primate eyes with human eyes enucleated and reoxygenated at different times after death, we show the exponential decay of the light response amplitudes has a time constant of 74 min. We find that both cause of death and donor age are useful parameters for predicting the recovery of retinal light responses in postmortem human macular tissue. Moreover, we present evidence that hypoxia and secondary acidosis, two modifiable factors, are primary contributors to the rapid loss of retinal light signaling after death. Finally, we show postmortem hypoxia rather than acidosis is the major cause of irreversible decay of the light response amplitudes. The criteria and methodology that we have established here for reviving electrical photoresponses in the postmortem human eye will serve as a new starting point for studying neurophysiology and disease in the human retina.

## Revival of light signals in the retina after death

Ischemic events leading to the cessation of blood circulation in the retina or brain rapidly leads to irreversible damage of the affected neurons [1-5]. Although there may be increased electrical activity, including the sensation of bright light soon after death (‘near death experience’) [6, 7], consciousness and brain activity is lost within ∼1 min of cerebral ischemia or cardiac arrest [6, 8-10]. To understand the timeline and cellular mechanisms blocking light-evoked retinal responses after death, we investigated how quickly retinal phototransduction and synaptic transmission are lost *in vivo* following circulatory death. We measured light-induced photoreceptor and ON bipolar cell responses using *in vivo* ERG in anesthetized mice before and after euthanasia by cervical dislocation (Fig. 1a, b). The photoreceptor component of the *in vivo* ERG signal was isolated in one eye by intravitreal injection of DL-AP4, an inhibitor of the metabotropic glutamate receptors (mGluRs). These experiments reveal a progressive reduction of both photoreceptor and bipolar cell driven ERG responses leading to almost complete loss of ERG activity within ∼5 minutes of death (Fig. 1c). Interestingly, the bipolar cell response amplitudes declined with time constant of 1.3 min, significantly faster than photoreceptor response (τ = 2.2 min, inset in Fig. 1c).

**Figure 1:**
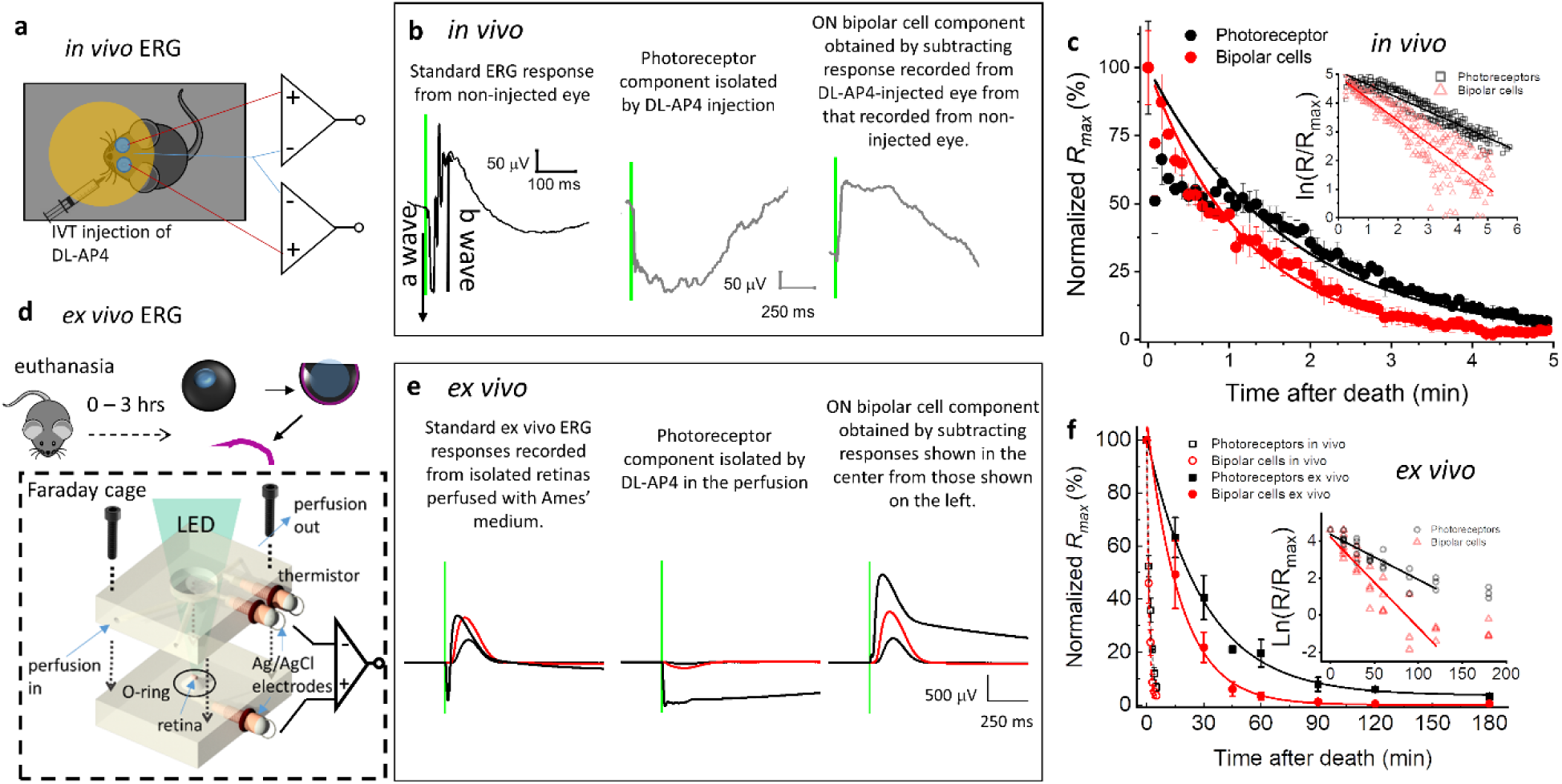
Postmortem retinal recordings and results. **(a)** Schematic of *in vivo* recordings illustrating isolation of photoreceptor responses using DL-AP4 intravitreal injections in one eye, example isolated photoreceptor responses shown in **(b)** (middle), which are subtracted from the standard ERG response from contralateral eye (left) to obtain a full ON-bipolar cell response (right). **(c)** responses to light flashes (10 Cd.s.m^-2^) obtained every 5 s during *in vivo ERG* experiments shown in (a), normalized to the first response taken. Inset shows *Ln* of normalized responses for all animals with a linear fit plotted for both bipolar cell and photoreceptor responses. **(d)** *Ex vivo* electroretinogram recordings from retinas isolated at different delays to enucleation, mounted onto custom built specimen holder, under constant perfusion of media. Responses are recorded via Ag/AgCl electrodes mounted above and below retina to different intensities of LED flashes. Example traces from experiments demonstrating the subtraction of isolated photoreceptor responses **(e, center)** from standard responses **(e, left)** to obtain an ON bipolar cell response **(e, right)**, red trace shows response to 26 photons µm^-2^. Mean responses from postmortem retinas with different enucleation delays normalized to maximal response size obtained from immediately enucleated eyes **(f)**. Inset shows Ln of normalized responses from all retinas with linear fits.

Although the transmission of light signals in the retina was quickly abolished after death *in vivo* as expected, several prior studies have shown that the electrical activity of individual neurons could be at least partially restored after extended times of complete cerebral ischemia [9, 11-14]. Thus, we asked if the activity of retinal neurons could be revived by restoring normal pH, the delivery of oxygen and select nutrients *ex vivo* (see Methods). To answer this question, we carried out *ex vivo* ERG experiments from isolated mouse retinas placed in a specialized specimen sholder (Fig. 1d) and supplemented with oxygenated and nutrient rich medium at different death to enucleation times, using freshly enucleated eyes as the control (Fig. 1e-f). Under these conditions, we measured significant recovery of photoreceptor and ON bipolar cell light responses (Fig. 1f). While the restored absolute amplitudes of photoreceptor and ON bipolar cell light responses declined with increasing death to enucleation times (Fig. 1c), small light responses could be measured from eyes enucleated even up to 3 hours after death (Fig. 1f). Although this was partly due to higher signal-to-noise ratio in the *ex vivo* ERG experiments, it is evident that the decay of restored retinal responses is significantly slower than the fast decline of the postmortem *in vivo* ERG light responses (Fig. 1f). Indeed, pharmacologically isolated *ex vivo* ERG photoreceptor and ON bipolar cell responses decayed >10-fold slower than those during postmortem *in vivo* recordings (Fig. 1f). Similar to our findings *in vivo*, ON bipolar cell responses decayed about two times faster with increasing enucleation delay than photoreceptors (see inset in Fig. 1f). These results demonstrate that even after all neuronal activity has ceased after death, significant light-evoked photoreceptor responses can be restored in retinal explants. Our results are consistent with prior studies that have reported light-evoked responses of photoreceptors[15-18], ganglion cells [19, 20] and rarely a residual ERG bipolar cell response (b wave) [15, 16] in isolated rat, pig and human peripheral retinas from eyes that had been enucleated up to several hours postmortem.

## Phototransduction after death

Having observed a gradual decline of the absolute amplitude of the light responses recovered *ex vivo* with increasing enucleation delays, we studied how death to enucleation times affects the light-sensitivity and kinetics of phototransduction recovery. Using *ex vivo* ERG, we first determined the light intensity required to generate a half-maximal response (*I*_*1/2*_) at different times after death and then compared dim light flash photoreceptor response amplitudes normalized to their respective *R*_*max*_ and flash intensity (*S*) (Extended Data Fig. 1a, d). Surprisingly, there were no significant differences in the *S* or *I*_*1/2*_ across all of the post-mortem delays measured, indicating that an individual photon will close the same fraction of the light-sensitive CNG channels in revived photoreceptors after different death to enucleation times as in photoreceptors before death.

While gain of phototransduction is not significantly affected by enucleation delay, it is possible phototransduction activation and inactivation reactions are affected. To assess the kinetics of the phototransduction activation and inactivation reactions we quantified the gain of the phototransduction activation reactions (*A*, s^-2^, see Eq. 1 in Methods) as well as time to peak (*t*_*p*_) and integration time (*T*_*i*_) of the dim flash responses respectively (Extended Data Fig. 1g-f). There is a trend towards smaller *A* up to 90 min postmortem, and the amplification constant at 90 mins is less than that of freshly enucleated eyes (Extended Data Fig. 1e). There is significant lengthening of *t*_*p*_ and *T*_*i*_ of the dim flash responses after 90 min postmortem, indicating deceleration of phototransduction inactivation reactions by extended enucleation delays (Extended Data Fig. 1f). These results suggest that phototransduction activation and inactivation reaction kinetics cannot be fully restored in the eyes that have been recovered more than ∼90 mins postmortem. However, in general the changes in the gain and kinetics of phototransduction are modest when compared to the significant and gradual decay of *R*_*max*_ after death (see Fig. 1f).

## Potential mechanisms for the loss of light signals after death

Our results above demonstrate an exponential decay of the amplitudes of photoreceptor and ON bipolar cell light responses recovered *ex vivo* after death (Fig. 1d). One interpretation is that photoreceptor and ON bipolar cells are undergoing apoptosis, with mean lifetimes of 41 and 20 min after death, respectively. To test this hypothesis, we visualized activated caspase-3 protein in sections from eyes enucleated at different times after death to determine if apoptosis was occurring. We detected only fractional numbers of activated caspase-3 positive cells in retinas from wild-type mice enucleated at 0, 60 or 180min after death (Fig. 2a) whereas activated-caspase 3 could be clearly detected in control retinas from 21 day old p23h mice (heterozygous), a well-characterized photoreceptor degeneration model (Fig. 2a, [21]). These results suggest that the loss of photoreceptor and bipolar cell function is not due to the induction of apoptosis within 3 hours of death.

**Figure 2.**
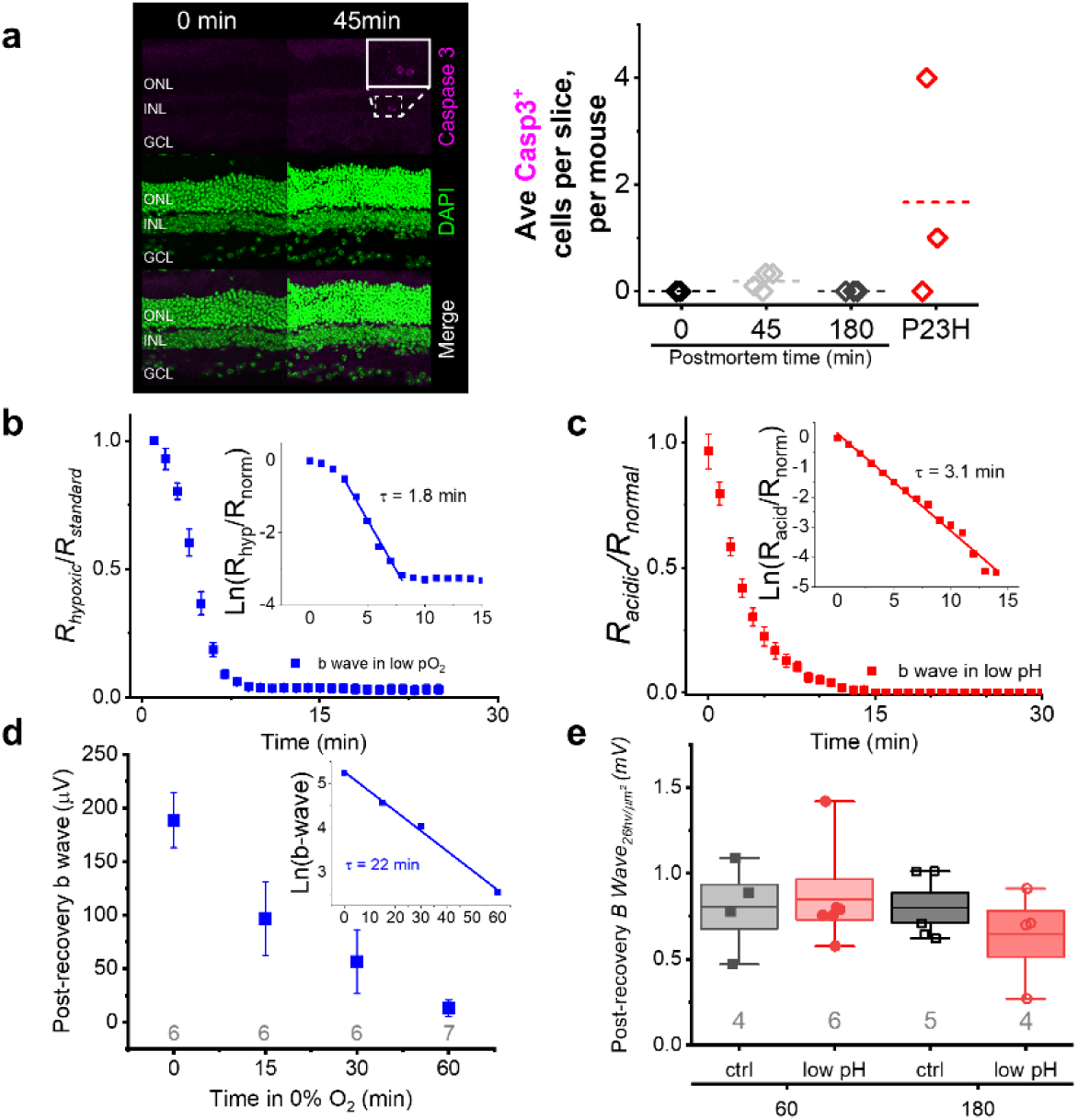
**(a)** Labeling of retinas at 0, 45- and 180 min postmortem with anti-activated caspase antibody, left shows representative control and postmortem retina. Box highlights two positively labelled retinal neurons. Plot shows the mean positive cells per slice, per mouse for each condition (dashed line), positive control values are from P14 *Rho*^*P23H/WT*^ mouse retinas. **(b)** Decline in ON-bipolar cell response amplitudes measured during perfusion in low oxygen, normalized to maximal amplitude obtained in normal oxygen concentration perfusion. Inset shows exponential fit to plotted Ln of hypoxic B-waves with rate of decay (τ). **(c)** Decay of ON-bipolar cell responses measured during perfusion in low pH (pH 6.8) Ames normalized to response amplitude obtained during normal pH Ames perfusion. Inset shows linear fit of the Ln of the normalized ON-bipolar cell responses, with a decay rate (τ) of 3.1 min. **(d)** ON-bipolar cell response recovery after incubation in low oxygen Ames for 0-60 min, inset shows the Ln of values and a rate of decay of response recovery of 22 min. **(e)** ON-bipolar cell response amplitudes to light stimulus of 26 photons µm^-2^ after incubation in low pH Ames for 60-180 min.

Retinal oxygen consumption is higher per gram than in any other tissue [22]. We hypothesized that hypoxia and/or acidification of the retinal tissue influences the postmortem decay and recovery of phototransduction and transmission of light signals in the mouse retina. To test this, we measured light responses of photoreceptor and ON bipolar cells from isolated retinas enucleated immediately after death (t=0) and perfused under low PO_2_ and increased H+. To maintain low oxygen, we used our closed *ex vivo* ERG system and an oxygen sensor to accurately control PO_2_ at the retina while recording ON bipolar cell responses to quantify their dependence on the availability of oxygen. We found that reducing PO_2_ from the standard level used in most electrophysiology studies (95% in the reservoir and 80% at the retina, see Methods) to ∼5% (or ∼40 mmHg) did not compromise phototransduction or transmission of light signals. However, reducing oxygen to 2.5%, below its expected value *in vivo* [23], quickly compromised light signaling in the retina (Fig. 2b). The logarithmic amplitude plot revealed essentially three phases of amplitude decay. The first decay phase is probably a result of the gradual decline in PO2 in the tissue after switching to the low oxygen perfusion. A linear fit to phase two, during which the majority of the decay occurred, revealed an exponential decay time constant of 1.8min (inset in Fig. 2b), slightly slower than observed for the postmortem decline of the bipolar cell response *in vivo* (Fig. 1c).

Next, we looked at the effect of lowering pH under standard nominal 95% PO_2_ conditions. Previous studies demonstrate pH declines to ∼pH 6.8 in postmortem eyes [24]. Based on this value, we compared photoreceptor and ON bipolar cell responses under normal (7.4) and low (6.8) pH by adjusting the level of bicarbonate in the perfusion medium (containing constant 5% CO_2_). As with very low PO_2_, exposure of the retina to low pH led to a rapid reduction of both ON bipolar and photoreceptor cell responses (Fig. 2c). Low pH exposure caused exponential decay of the b wave amplitudes with time constant of 3.1min. These results suggest that postmortem acidification and hypoxia would both lead to a rapid decay of retinal light signaling after death *in vivo* as we demonstrated in Fig. 1a-c.

We next asked whether extended postmortem exposure to low PO_2_ or higher [H^+^] leads to irreversible changes that could explain the inability to recover full bipolar cell response amplitudes with increasing enucleation delays (Fig. 1f). To answer this question, we incubated mouse eyes in low PO_2_ (2.5%) or low pH (6.8) for up to 180 min and measured *ex vivo* ERG ON bipolar cell response amplitudes from retinas dissected from these eyes under standard conditions (95% PO_2_ and pH 7.4). Hypoxic incubation had an irreversible effect on the amplitude of responses even after a recovery period in normal perfusion conditions (Fig. 2e). Interestingly, the recoverable bipolar cell response amplitude decayed according to a single exponential function (τ = 22 min) with increasing exposures to hypoxia following kinetics strikingly similar to that observed in our *ex vivo* ERG postmortem data (τ = 20min, inset in Fig. 1f, red). On the other hand, while loss of b-wave amplitude in low pH occurred relatively rapidly (Fig. 2c), bipolar cell function can be recovered after 180 min incubation in low pH (Fig. 2d). These results indicate that postmortem hypoxia rather than acidosis is the rate-limiting factor explaining the incomplete restoration of retinal light responses after extended death to enucleation times.

To further investigate factors that may contribute to rapid loss of synaptic transmission from photoreceptors to ON bipolar cells after death, we used Computational Molecular Phenotyping (CMP) [25] to quantify changes in the metabolic profiles of the retinas. We find that mouse retinas from eyes enucleated at 45 min postmortem have several key changes in their molecular profiles compared to control retinas fixed immediately after euthanasia. We quantified these differences in three cell types: photoreceptors (outer segments and somata), bipolar cells, and Müller cells. Taurine concentration is reduced in all three cell-subtypes after 45min postmortem. Both glutamine and glutamate are reduced in bipolar cells, photoreceptor cell somata, outer segments, and Müller glia at 45 min post-mortem (Extended Data Fig. 3). The loss of both the neurotransmitter required for transmission of photoreceptor potentials to the bipolar cells, as well as the loss of the key metabolite involved in the production of this neurotransmitter, could contribute to the rapid loss of photoreceptor-derived light response of ON bipolar cells after death.

## Recovery of phototransduction in the human macula after death

We demonstrated that limited recovery of photoreceptor and ON bipolar cell light responses is possible several hours after death in the rod-dominated, nocturnal mouse. However, humans are diurnal with a high cone-density in the central fovea/macula region of the retina. Thus, we next tried to recover light responses from macular cone photoreceptors in human donor eyes collected 0.5 – 5 hours postmortem. Prior studies have reported light responses of rods and cones in human peripheral *ex vivo* retina samples from eyes removed from live patients (due to cancer or retinal surgery, [26, 27]), or up to several hours postmortem from organ/research donors [15-17]. However, to the best of our knowledge, light responses of human cones in the macula have not been reported. We optimized our *ex vivo* ERG method to record photoreceptor responses from a 5 mm (diameter) patch of the retina centered approximately at the fovea in macaque eyes enucleated and placed into oxygenated Ames’ media within 10 minutes after euthanasia (see Methods for details). A representative photoreceptor response family (isolated using DL-AP4) to light flashes ranging from ∼200 to 185,000 photons μm^-2^ is shown in Fig. 3a. These responses are significantly larger than cone responses from peripheral samples obtained using a rod-saturating pre-flash or background light (Extended Data Fig. 4) probably due to higher cone density in the macula compared to peripheral primate retina. To obtain human eyes with viable retinas, Eye Bank recovery personnel were provided with Ames medium in light-tight containers. Eyes were recovered within 5 hours from death and placed immediately in Ames medium on-site. Following the enucleation, the eyes were delivered in Ames’ medium in darkness to our laboratory within an hour. As detailed in Methods we first dark-adapted eyes in oxygenated Ames’ followed by dissection of a 5mm patch of the macula as described for macaque eyes above. Using these protocols, we were able to record first *ex vivo* light responses from the human macula (Fig. 3b and c). The amplitudes of light responses in the macula from an eye enucleated an hour postmortem were similar to those recorded from a freshly enucleated macaque. Although response amplitudes were smaller in the macula obtained from an eye harvested >2 hours after death, we were also able to record robust light responses from the maculae of human eyes harvested up to 5 hours postmortem (Fig. 3d).

**Figure 3.**
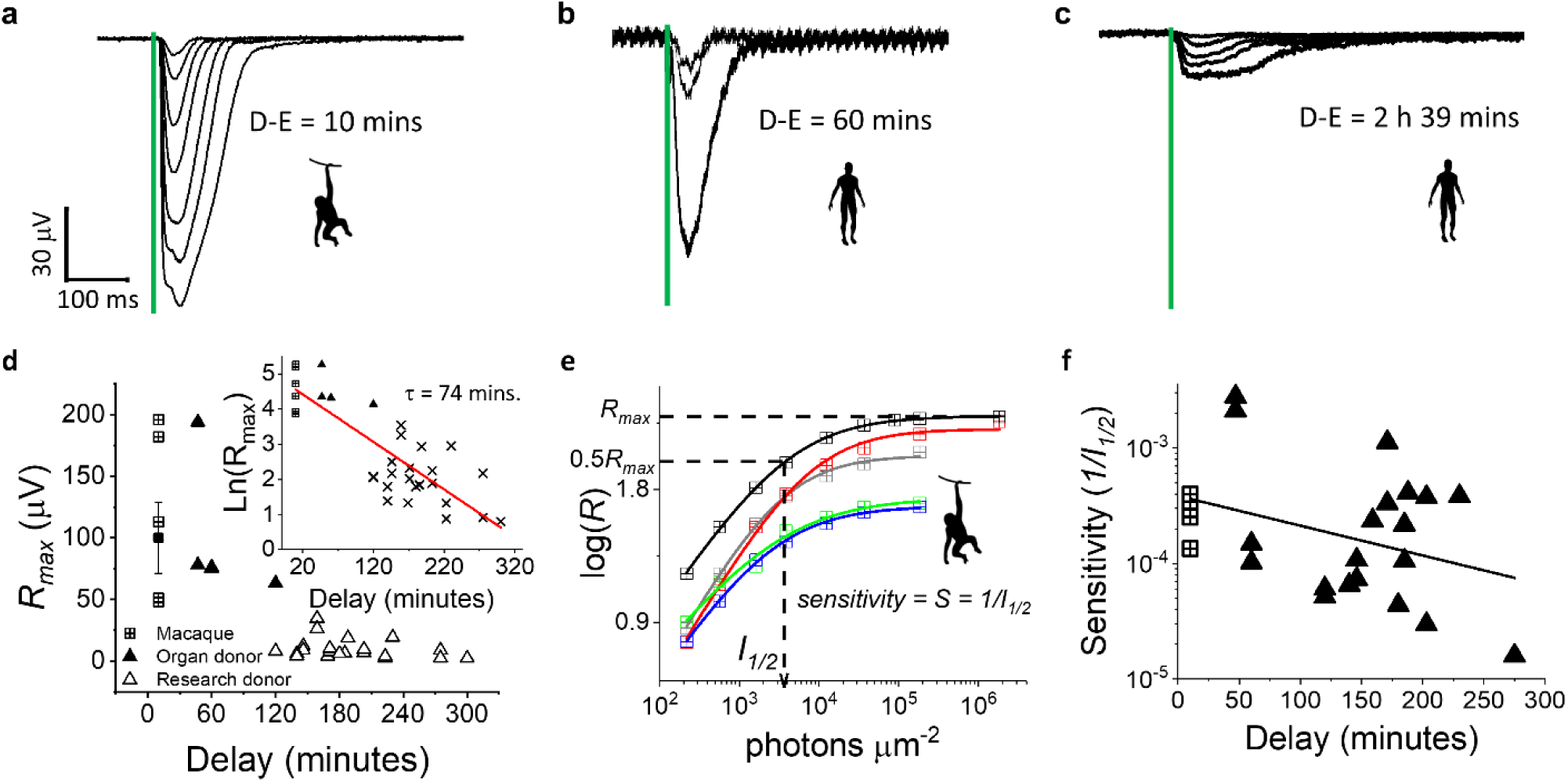
Light responses of macular cones from freshly enucleated (10min death to enucleation delay, D-E) macaque **(a)** human organ donor **(b)** and research donor **(c)** eyes, timing of light stimulus indicated with green line. **(d)** The maximal response obtained from maculae from each tissue type plotted against enucleation delay. Inset shows Ln of *Rmax* from each macula, and linear fit in red with a tau of 74 min. **(e)** Example plot demonstrating how sensitivity, S (1/I1/2) was calculated. Responses from maculae to different light intensities were plotted and a Hill curve fitted to determine the intensity required to produce a half-maximal response size (Eq. 3 in Methods). **(f)** Sensitivities of each macula plotted against enucleation delay, linear fit in black (no significant correlation between delay and S).

Experiments using human donor eyes introduced several uncontrolled variables not present in our macaque experiments, including postmortem enucleation delay, cause of death, co-morbidities, retinal disease, and age of the donors. We collected data from 20 individual human donors to better understand how these parameters may contribute to the viability of macular cones. Our first hypothesis was that enucleation delay would be the most critical factor in determining the response amplitudes and light sensitivity of human macular cones. To test this hypothesis, we plotted the *R*_*max*_ of our macaque experiments together with our human donor eye experiments as a function of postmortem enucleation delay (Fig. 3d). We fitted a single exponential function to the *R*_*max*_ data (red trace in Fig. 3d) and performed Pearson’s correlation analysis between Ln(*R*_*max*_) and enucleation delay (see inset in Fig. 3d). As expected from our mouse data, these analyses indicate that there is a highly significant and strong negative correlation between *R*_*max*_ and enucleation delay, and that the response amplitudes of primate macular cones decay with a single exponential time constant of 74min, almost two times longer than that determined for the mouse photoreceptors (Fig. 1f). To test if this is rod vs. cone or species difference, we first determined the effect of postmortem delay on cone photoreceptor responses using *Gnat1*^*-/-*^ mice. We found that the time constant of decay is somewhat longer (59min, see Extended Data Fig. 2a) for mouse cones compared to mouse rods (41 min, inset in Fig. 1f, black), and although still appears to be shorter than that for the human macular cones there was no statistically significant difference between mouse and human cones.

We also compared sensitivity of human macular cones to that of freshly enucleated macaque eyes and found no significant correlation between sensitivity and enucleation delay (Fig. 3f). This result is consistent with our mouse data where rod or cone sensitivity, measured as the intensity required to generate a half-maximal response (*I*_*1/2*_), was not affected by the postmortem enucleation delay (Extended Data Fig. 1c, and 2b).

Although we were able to record photoreceptor responses from human donor eyes up to 5 hours after death, we did not observe post-receptor components of the *ex vivo* ERG signal in any of the human macula samples (n = 17). In this study we did not systematically study rod or cone light responses in human peripheral samples but recorded a positive ERG b-wave, indicative of post-receptoral activity, only from peripheral samples of one donor (see Extended Data Fig. 5). Previous studies have also observed *ex vivo* ERG b waves on occasion from postmortem peripheral human retina samples [15, 16]. Although lack of observable ON bipolar cell light responses is consistent with faster decay of the mouse ON bipolar than photoreceptor cell responses as well as prior studies showing that b wave is extremely sensitive to ischemia/anoxia [28, 29], the complete lack of ERG signal from bipolar cells in donor human maculae is surprising since we consistently measured residual rod and cone ON bipolar cell responses from mice up to 3 hours after death (albeit they were extremely small). Thus, future experiments are needed to understand factors contributing to the lack of robust ERG b waves in postmortem human eye tissue.

Since the decay of *R*_*max*_ as a function of enucleation delay is exponential, the difference in absolute amplitudes is relatively small in the group of donors where enucleation delay is between 2 and 5 hours. We used this group of donors which included 17 individuals between 53 and 86 years of age (Fig. 4a, mean age 67 ± 2), and died due to sepsis, acute cardiac event (ACE) or stroke (Fig. 4b, n = 7 donors, n = 2 donors, and n = 2 donors respectively) to study the effects of various cause of death parameters on *R*_*max*_ of the human macular cones. Comparing the *R*_*max*_ for the three most common causes of death for our donors demonstrates that sepsis produced a significantly lower *R*_*max*_ when compared to stroke or ACE, and that ACE yielded significantly lower *R*_*max*_ than stroke (Fig. 4c). We next tested if *R*_*max*_ is affected by donor age by plotting the maximal responses obtained from each macula as a function of the age. Surprisingly, we find that there is a significant positive correlation with *R*_*max*_ and age, i.e. maculae were more viable from older patients (Fig. 4d). A possible explanation for the larger responses obtained from older donors could either be related to our criteria to exclude donors with ocular disease history, or to the possibility that more liquid vitreous in older patients promotes better access of oxygen and nutrients to the retina as well as easier sample preparation.

**Figure 4.**
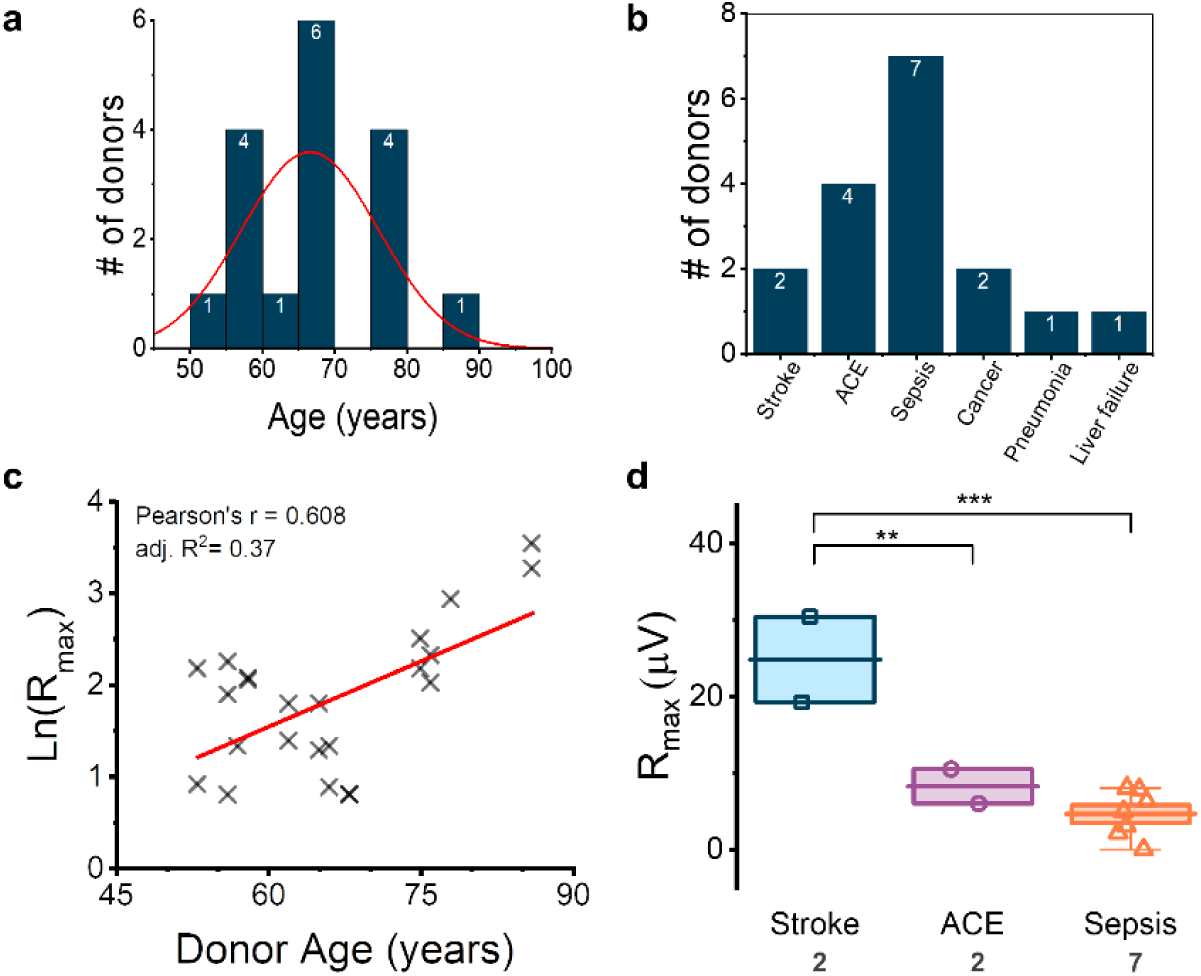
**(a)** Histogram of research donor ages, with numbers of donors represented within the bars. **(b)** Histogram of numbers of donors obtained and their primary cause of death as noted on tissue donor forms. ACE, acute cardiac event. **(c)** The maximal response obtained from each donor eye plotted against donor age, with linear fit showing small positive correlation (Pearson’s r = 0.606, R^2^ = 0.37) **(d)** Average maximal amplitude obtained from each research donor according to primary cause of death, data uses average of responses obtained from both maculae where responses could be obtained recorded from both donor eyes. Significance was tested using One-way ANOVA with Holm-Bonferroni means comparisons, ** indicates p < 0.025, *** indicates p< 0.01667. Boxes indicate SEM, central line is mean value, and whiskers show minimum to maximum values. Individual data points are overlaid, n numbers (donors) are indicated below cause of death.

In summary, we have demonstrated the first successful recordings of human cone photoresponses in the macula from donor eyes enucleated up to 5 hours postmortem. We have identified important factors that affect the quality of the responses from these postmortem eyes including the cause of death and enucleation delay. We also present evidence that postmortem hypoxia and secondary acidosis contribute to the loss of light signaling after death. Interestingly, whereas the effects of acidosis were largely reversible, hypoxia caused a relatively quick irreversible loss of signal transmission in the retina. The methods and criteria described in this paper will serve as a starting point for future studies to understand human high-acuity color vision mediated by the macular cone photoreceptors and how it is affected in aging and disease.

## Methods

### Animal Handling and ethical statement

*C57Bl6/J*, and *Gnat1*^*-/-*^ [30] mice were bred in house or purchased at 8-10 weeks of age from Jackson Laboratory (Bar Harbor, ME, USA). All mice were housed within the University of Utah’s animal facilities. Animals were maintained on a light dark cycle of 12 h light/12 h dark. All experiments were approved by the Institutional Animal Care and Use Committee of the University of Utah and conducted according to the ARVO Statement for the Use of Animals in Ophthalmic and Vision Research. Unless otherwise noted, all animals used were between 6-20 weeks old, and both males and females used for experiments. Macaque (*Macaca fascicularis*) eyes were kindly provided by Dr. Chad Faulkner and Dr. Joel Perlmutter at Washington University in St. Louis. Care of macaques were in accordance with Guide for the Care and Use of Research Animals and all experimental protocols were approved by the Washington University School of Medicine Animal Studies Committee.

### *In-vivo* ERG

We assessed light-evoked electrical activity of photoreceptors and ON bipolar cells using *in vivo* electroretinogram (ERG) in wildtype C57Bl/6J mice. Wildtype mice were fully dark adapted overnight and anesthetized for 20 minutes with 2% isoflurane in room air using a SomnoSuite Low Flow Anesthesia System (Kent Scientific, Torrington, CT, USA). Pupils were dilated with 2% atropine sulfate eye drops (Akorn Pharmaceuticals, Lake Forest, IL, USA) and corneas were lubricated with a mixture of phosphate buffered saline (PBS) with Goniovisc (2.5% hypromellose solution, Contacare, Gujarat, India). Euthanasia was by cervical dislocation and recording was started within 30 seconds of death. ERG waveforms were recorded using corneal contact lens electrodes, a subdermal needle electrode inserted between the eyes was the reference electrode, a subdermal needle at the base of the tail as the ground electrode. Responses to flash intensities of 0.5 cd.s.m^-2^ every 5 seconds (365nm ± 9nm wavelength) were collected using a ColorDome (Diagnosys LLC, Lowell, MA, USA) and Espion V6 software.

### *Ex-vivo* ERG

*C57BL/6J*, and *Gnat1*^*-/-*^ mice were dark adapted overnight (8-12h), before euthanasia by CO_2_ inhalation and cervical dislocation. All procedures after dark adaptation were performed under dim red light. Eyes were enucleated into freshly prepared Ames media (#A1420, Sigma-Aldrich, St Louis, MO, USA) bubbled with 5 % CO_2_/95% O_2_ unless otherwise stated. Eyes were hemisected, lens and cornea removed, and retina isolated from the eyecup.

*Ex vivo* Electroretinogram (ERG) recordings were performed as described previously [31]. Isolated retinas were mounted photoreceptor side up onto the specimen holder and perfused with Ames supplemented with BaCl_2_ (100µM, # 342920, Sigma-Aldrich, St Louis, MO, USA) to isolate inner retinal signaling (B-waves) and DL-2-Amino-4-phosphonobutyric acid (DL-AP4, 40µM, Tocris Bioscience, Bristol, U.K.) to isolate the photoreceptor component of the ERG signal (A-wave). The rate of perfusion was ∼5 ml/min, perfused media was temperature controlled at 37^°^C and bubbled with a 95% O_2_ and 5% CO_2_ mix.

ERG signal was first amplified (100X) and low-pass filtered at 300 Hz by a differential amplifier (DP-311, Warner Instruments, Hamden, CT, USA), and data further amplified (10X) and acquired at 10KHz using an integrated Sutter IPA amplifier/digitizer (IPA, Sutter Instrument, CA, USA). Light stimuli were provided either from a High-Power LED light source (Solis-3C, Thorlabs, Newton, NJ), with filter for green light and LED driver (DC2200, Thorlabs, Newton, NJ, USA), or a High-Power LED light source (LED4D067, 530nm, Thorlabs, Newton, NJ, USA). Light stimulus durations ranged from 2-20ms. The SutterPatch software (SutterPatch v2.0.2, Sutter Instrument, CA, USA) drove both stimulus generation and data acquisition via the IPA amplifier’s analogue output and input, respectively. Light stimuli were calibrated before experiments using a calibrated photodiode (FDS100-CAL, Thorlabs, Newton, NJ, USA) and flash intensities converted to photons µm^-2^.

### Effect of post-mortem to enucleation time on photoreceptor and ON-bipolar cell function

For experiments simulating the effect of delay to enucleation to the ERG, mice were euthanized as above. Enucleation was delayed by 15 to 180min, or control eyes enucleated immediately after euthanasia. *Ex vivo* ERG was then carried out using either Ames + BaCl_2_ to measure bipolar cell responses with photoreceptor responses, or Ames + BaCl_2_ + DL-AP4 to measure photoreceptor responses in isolation. Subtraction of the pharmacologically isolated photoreceptor responses from the responses of the photoreceptors combined with bipolar cells provides the amplitude of the bipolar cell response (Figure 1e).

### Effect of low pH and low oxygen on bipolar cell function

For experiments simulating the effect of postmortem hypoxia we compared light responses between standard conditions where the perfusion solution was continuously bubbled with standard 95% O_2_/5% CO_2_ gas mixture to one where most of the oxygen was replaced by nitrogen (N_2_). When we saturated our Ames’ medium with 95% O_2_/5% CO_2_, 80% PO_2_ was measured with an oxygen microelectrode at the retina, whereas when perfusing with 95% N_2_/5% CO_2_ the PO_2_ dropped down to 2.5% at the retina.

For experiments comparing the effect of low pH on ON-bipolar cell function, we titrated the concentration of sodium bicarbonate in the Ames perfusion solution used until a pH of ∼6.8 was achieved under the conditions maintained during incubation or perfusion. In both conditions, retinas were perfused with the modified (low pH or low oxygen) or control Ames solutions, and responses to a stimulus 90 photons µm^-2^ taken over the course of 60 minutes.

To understand if changes in the ON-bipolar cell function is reversible, we incubated the entire eye in Ames of either low pH or low O_2_, obtained as described above, between 15min to 3 hours, with control eyes incubated for the same period in pH 7.4 Ames bubbled with the regular gas (95% O_2_/5% CO_2_) mixture. Since eyes for low oxygen experiments were bathed in Ames medium saturated continuously with 95% N_2_/5% CO_2_ in a closed container, we expect that they were exposed to 0% O_2_ for the duration of the incubation. Following this initial incubation in low pH/low O_2_, or control conditions, the retinas were dissected, mounted in the *ex-vivo* ERG specimen holder, and perfused for 20 minutes (low O_2_ experiments) or one hour (low pH experiments) with warmed Ames bubbled with 95% O_2_/5% CO_2_. Following this period of perfusion in normal Ames, we took a response profile from the retina to measure the recovery of responses.

### Human Organ Donor and Macaque *ex vivo* ERG

Enucleated eyes from three individual macaques (*Macaca fascicularis*), that were part of other research projects, were obtained after their scheduled euthanasia (with pentobarbital, >150mg/kg IV). *Ex vivo* ERG experiments were conducted in the laboratory that was ∼10 min walking distance from the location of euthanasia. Macaques were not dark adapted prior to euthanasia and bright surgical lights were used during eye enucleation. Thus, to promote pigment regeneration and dark adaptation of photoreceptors we incubated the whole eyeballs in oxygenated Ames’ media for 20 – 30 minutes in darkness. This may not have been enough to ensure a complete dark adaptation of rod photoreceptors but was a compromise to minimize total postmortem time and to support cone dominance in the *ex vivo* ERG signal. After incubation, cornea, lens, and most of the vitreous was removed before a 5 mm5mm patch of the retina was taken centered at the fovea, visible as a darker spot 3 mm temporal from the optic nerve. These macular patches were placed on a custom-build *ex vivo* ERG specimen holder with a 4mm effective recording area. Light responses to a homogenous 5mm spot of light covering the whole recording area were recorded from 3 individual macaques and 5 maculae.

### Human Research Donor Eye *ex vivo* ERG experiments

Donor eyes obtained from the Utah Lions Eye Bank (n = 17 pairs) between 2-5h post-mortem and from San Diego Eye Bank (n = 3 pairs) 45min – 2h post-mortem, after full consent was given by family members and in accordance with the Declaration of Helsinki. All retinal tissues and data were de-identified in accordance with HIPAA Privacy Rules. Exclusion criteria given to the eye bank included diagnoses with ocular disorders such as AMD and diabetes. Whole globes were stored in bicarbonate buffered Ames in the dark after enucleation and during transport and allowed to dark-adapt a further 30-60min (depending on transport time). The lens and cornea were removed from the eye, and a 5mm biopsy punch was used to take tissue punches from either Macula or Peripheral retina, neural retina was then separated from the underlying retinal pigment epithelium (RPE) and choroid. Each retinal punch was mounted in a custom-built specimen holder and perfused with Ames containing 100µM BaCl_2_ bubbled with 95% O_2_ and 5% CO_2_ and warmed to 37°C during experiments.

### Immunohistochemistry of the post-mortem retina

Wildtype mice were euthanized according to IACUC guidelines. Control eyes were enucleated in the dark immediately or delayed for 45, or 180 minutes before enucleation. All eyes were then fixed for 10min dark in 4% PFA with 30% glucose PBS, the cornea was punctured, and a second 10-minute fixation was carried out. Eyes were washed in PBS and the cornea and lens removed. Eyecups were cryoprotected for 30 minutes at room temperature in 30% sucrose PBS, before incubation in 30% sucrose PBS overnight at 4°C. Eyecups were then embedded in OCT medium (Tissue-Tek, Sakura Finetek, Torrance, CA, USA) before slicing 12µm sections and mounting on slides. To minimize differences in staining that could be attributed to sample preparation, eyecups from two separate treatment groups were mounted on each slide. Slides were blocked with 10% normal goat serum (NGS) in 0.1% PBS-Triton-X for one hour at room temperature, before incubating with Anti-activated Caspase-3 antibody (catalog# 559565, 1:500 dilution, BD Pharmingen, San Jose, CA, USA) in blocking buffer overnight at 4°C. Slides were washed in PBS before incubating in secondary antibody (Goat anti Rabbit Alexa 594, 1:1000, Invitrogen, Carlsbad, CA, USA) at room temperature for one hour. Slides were then mounted with a coverslip and Fluoromount-G mounting medium with DAPI (Invitrogen, Catalog# 00-4959-52, Invitrogen, Carlsbad, CA, USA). Images were acquired on an Olympus confocal (FV1000, Olympus Scientific Solutions, Waltham MA, USA), with a 40X oil immersion objective (Olympus UAPO 40X oil immersion, NA 1.35, Olympus Scientific Solutions, Waltham MA, USA) at 640 x 640 pixel resolution (pixel size = 0.497µm^2^). Imaging settings were optimized to the fluorescence signal on a positive control slide (P23H heterozygous retina), and then maintained for the rest of the slides. A single image was taken from several sections from each eye in each condition, and the number of activated-caspase 3 labelled cells in each retinal section averaged for each mouse, which was then used to calculate an average for each condition.

### Computational Molecular Phenotyping (CMP)

C57Bl/6J mice of 2 months of age of both sexes were euthanized by CO_2_ according to IACUC guidelines, and either decapitated and enucleated immediately (control) or after a 45min post-mortem delay. Methods are described in [25], briefly: Eyes were enucleated and fixed in buffered 2.5% glutaraldehyde/1% formaldehyde overnight, before resin embedding and serial sectioning. The sections were probed with IgGs for small molecules (see Table 1 for list) including: glutamate (E), glutamine (Q), and taurine (TT). Subsequent visualization was with secondary antibodies conjugated to 1.4nm gold (Cat. no. 2300, Nanoprobes, Yaphank, NY, USA), followed by silver intensification.

**Table 1.**
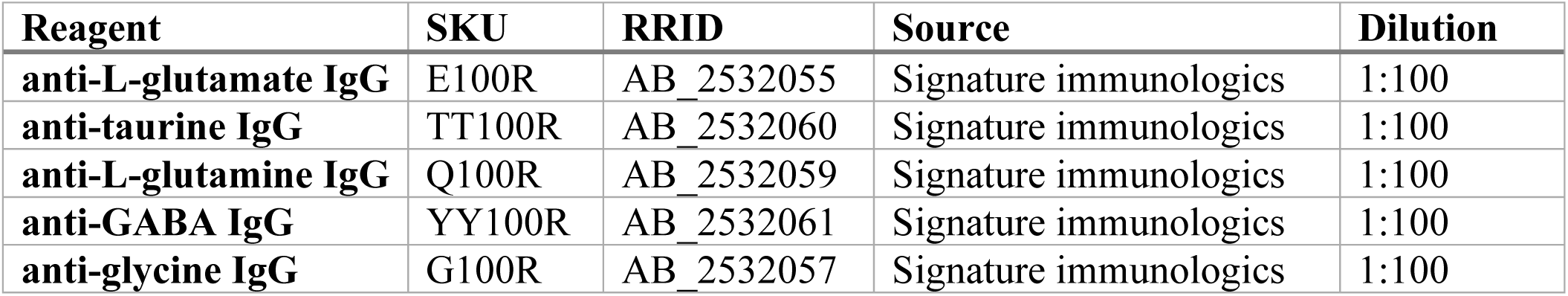
CMP reagents

Images were captured as 8-bit high resolution (243nm/pixel) images and registered with ir-tweak (https://www.sci.utah.edu/download/ncrtoolset.html) into image databases. Adobe Photoshop CC (Adobe, San Jose, CA, USA) was used for final image generation. For display only, raw data channels are linearly contrast-stretched. Molecular signals were visualized as selected RGB maps encoding Taurine, L-glutamine and L-glutamate to red, green, and blue color channels. Masks were created to isolate bipolar cell bodies, Muller cells in the outer nuclear layer, photoreceptor cell bodies and photoreceptor outer segments, to prevent contamination of signal from other cell populations. Monochrome images are density mapped. Histograms were taken from an area of retina spanning 200µm wide, for each retina sampled.

### Data Analysis

Data analysis, including statistical analysis and figure preparation in Origin 2018 (OriginLab, MA, USA), and edited in Adobe Illustrator CC 2019 (Adobe, San Jose, CA, USA). All data presented as mean ± SEM, and number of retinas in each experiment are indicated in figure legends. Normalized responses are calculated for each retina by dividing the response amplitude data by the maximal amplitude measured at the peak/plateau of the response to the brightest flash (R/R_max_). CMP images were prepared in Adobe Photoshop CC 2019 (Adobe, San Jose, CA, USA), and histograms of pixel intensities measured in FIJI [32].

### Statistics

Statistical differences between different experimental groups were typically analyzed by a one-way ANOVA, or two-way ANOVA with Holm-Bonferroni means comparisons test where appropriate. Comparisons of linear fits were carried out with the F-test, with p<0.05 considered a significantly different fit. Data are presented as mean±SEM. A value of *p*<0.05 is considered significant. N numbers are noted in the figure legends and on bar graphs and box charts.

### Equations

To determine the amplification constant for the dim responses of both rod and cones in the post-mortem retinas, we fitted to dim responses from both C57Bl6J and Gnat1^-/-^ mouse retinas a model developed by Lamb and Pugh to describe the kinetics of phototransduction activation [33]. The model may not accurately describe the kinetics of cone responses [34] but was used here to quantify and compare activation kinetics between immediate and post-mortem delayed cones. Here, *t*_*d*_ is a small delay corresponding to the small delay in recording of responses, as well as delays in phototransduction, and *A* is the amplification in s^-2^. Both *t*_*d*_ and *A* were allowed to vary to provide the best fit to the early rising phase of a dim flash response. Only intensity (F) was fixed to the intensity of light used (for C57Bl/6J retinas: 27 photons µm^-2^, for *Gnat1*^*-/-*^ retinas: 1010 photons µm^-2^).

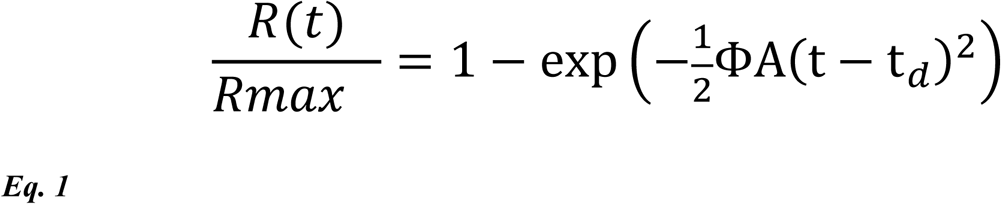

A Naka-Rushton function (Eq.2) was fitted to response amplitude data to determine flash strength producing 50% of R_max_ (I_1/2_):

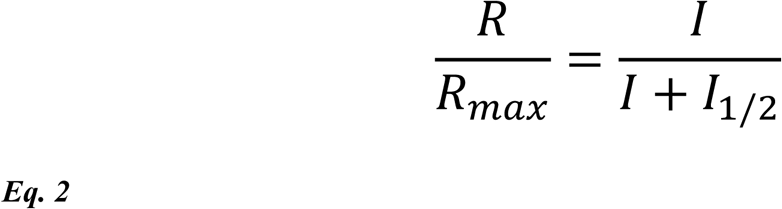

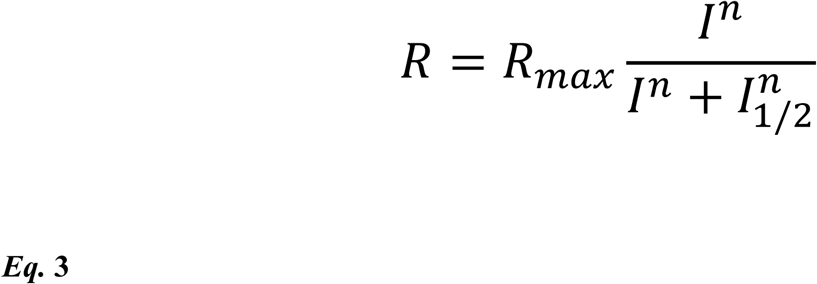

## Materials

## Acknowledgments

This work was supported by National Institutes of Health (P30 EY014800), and an Unrestricted Grant from Research to Prevent Blindness, New York, NY, to the Department of Ophthalmology & Visual Sciences, University of Utah. FV is supported by NIH grant EY026651, Research to Prevent Blindness / Dr. H. James and Carole Free Career Development Award, Diabetes Research Connection and International Retinal Research Foundation. BWJ is supported by the National Institutes of Health (R01 EY015128, R01 EY028927, P30 EY014800) and an unrestricted Research Grant from Research to Prevent Blindness, New York, NY to the Department of Ophthalmology and Visual Sciences, University of Utah. SP is supported by the National Institutes of Health (R01s: CA236352, DK115214, DK118278), Department of Defense (W81XWH1810645) and Department of Homeland Security (EMW-2016-FP-00788). AH is supported by Directed Research Philanthropic Funding. Authors thank Teemu Turunen for help with illustration of the *ex vivo* ERG specimen holder, and Dr. Chad Faulkner and Dr. Joel Perlmutter from Washington University in St. Louis for providing non-human primate eyes. Many thanks to the Utah Lions and San Diego Eye banks as well as LifeSharing for providing human donor eyes. We are also deeply grateful to those who donated their eyes and their relatives.

## Extended data figures

**Extended data Figure 1:**
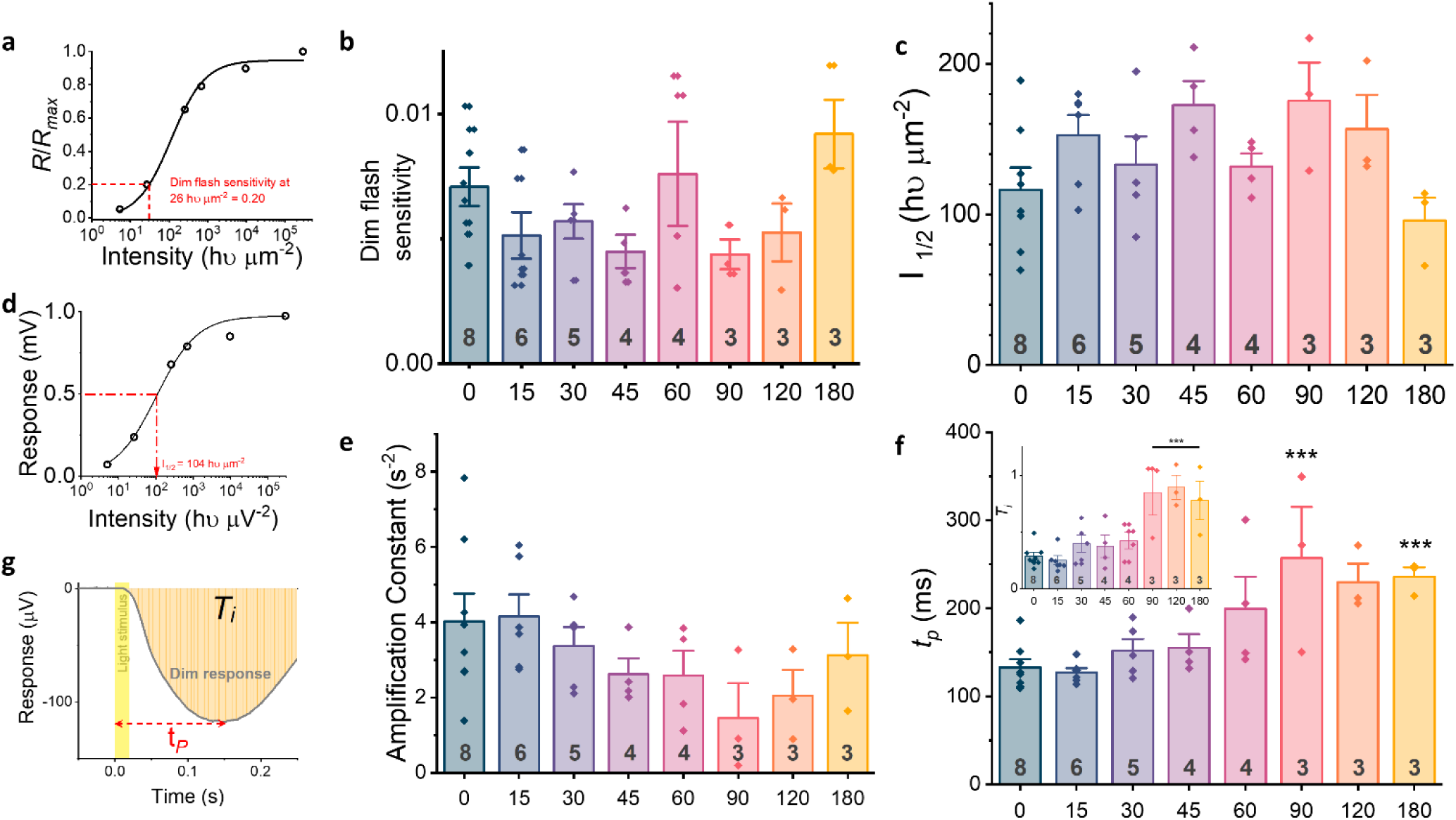
Example plot showing how dim flash response sensitivity is calculated **(a)**. Responses normalized to maximal response of retina plotted against intensity of light stimulus, curve fits Naka-Rushton function (Eq. 2) to amplitude data, dim flash sensitivity is defined as sensitivity at intensity producing ∼20% of the maximal response amplitude. Dim flash sensitivities of retinas from wildtype mouse retinas with different enucleation delays **(b)**, and I values from retinas of wildtype mice with different enucleation delays **(c)**. Example response plot showing how I_1/2_ was obtained **(d)**, responses of retina to each intensity presented is plotted and a Naka-Rushton fit applied to the plot. I_1/2_ is the intensity required to produce half the maximal response size. Amplification constant plotted against enucleation delay for each retina **(e)**. Time to peak (*t*_*p*_) and integral (T_i_, inset) of dim responses for each retina plotted against the enucleation delay **(f)**. (*** p < 0.001 vs 0min, 15min, tp) (*** p < 0.001 vs 0min, T_i_) Example dim flash response showing how time to peak (t_p_) and integral (T_i_) was calculated **(g)**. Time to peak is the time from light stimulus onset to the peak of the dim flash response. Integral (T_i_) is the area under the dim flash response curve (orange shaded area), divided by the response amplitude. Light stimulus is indicated as shaded yellow area. For all plots n is number of retinas and is indicated in dark grey at base of each bar, bar height indicates mean, error bars represent ± SEM. All comparisons are one-way ANOVA with Holm-Bonferroni means comparisons; p values are indicated in the legend for each graph.

**Extended data Figure 2:**
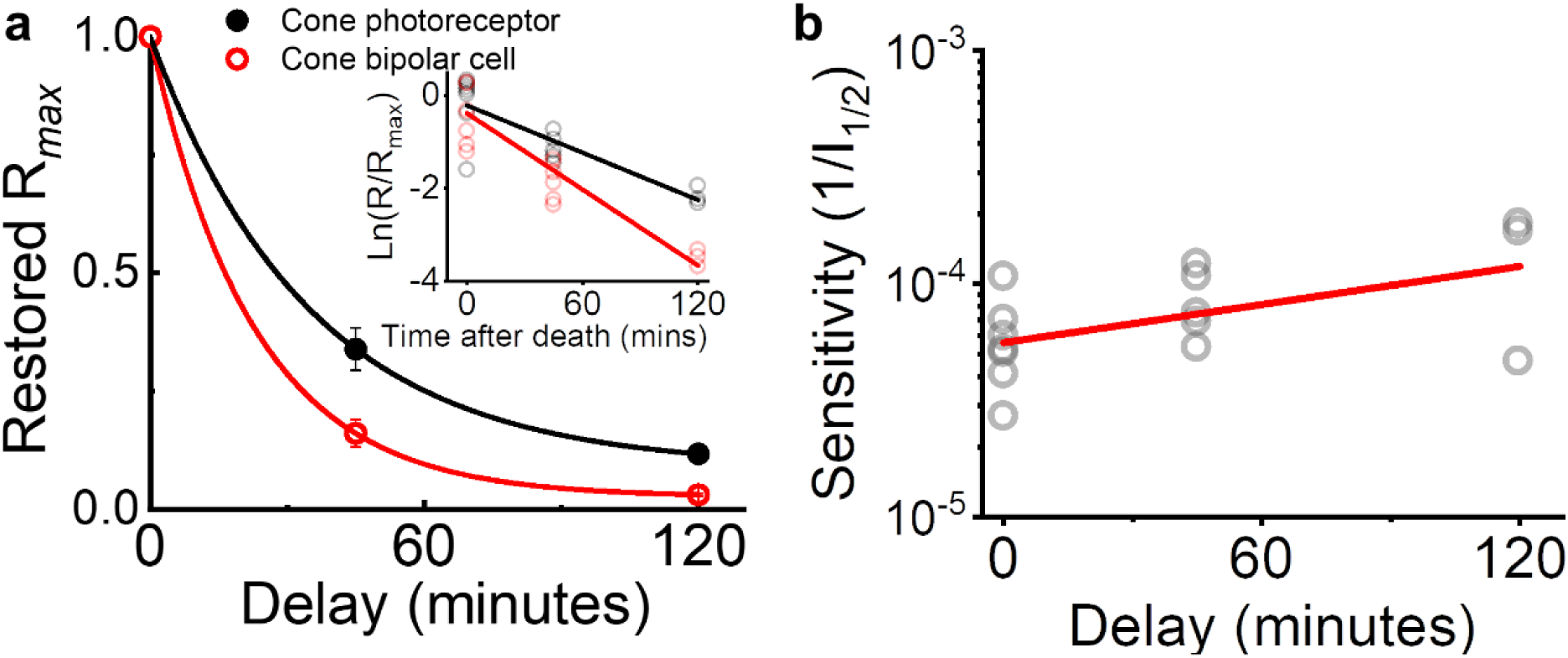
**(a)** The R_max_ of photoreceptors (black, filled) and bipolar cells (red, hollow) plotted as a function of enucleation delay, normalized to maximal response obtained with 0min enucleation delay. Smooth traces plot exponential decay function fitted to mean data. Inset plots logarithmic amplitude data for individual retinas, and linear fit (Photoreceptor fit: Pearson’s r = −0.851 with adj. R^2^ = 0.704, and τ = 59 min, Bipolar cell fit: Pearson’s r = 0.905 adj R^2^ = 0.819, and τ = 37 min). Sensitivity of retinas at different enucleation delays (**b**, 1/I_1/2_), Pearson’s r = 0.521 with adj. R^2^ = 0.215. For all plots, n (retinas) is 0min = 7, 45min = 5, 120min = 3, error bars represent ± SEM. Responses from individual retinas are plotted as hollow markers.

**Extended data Figure 3.**
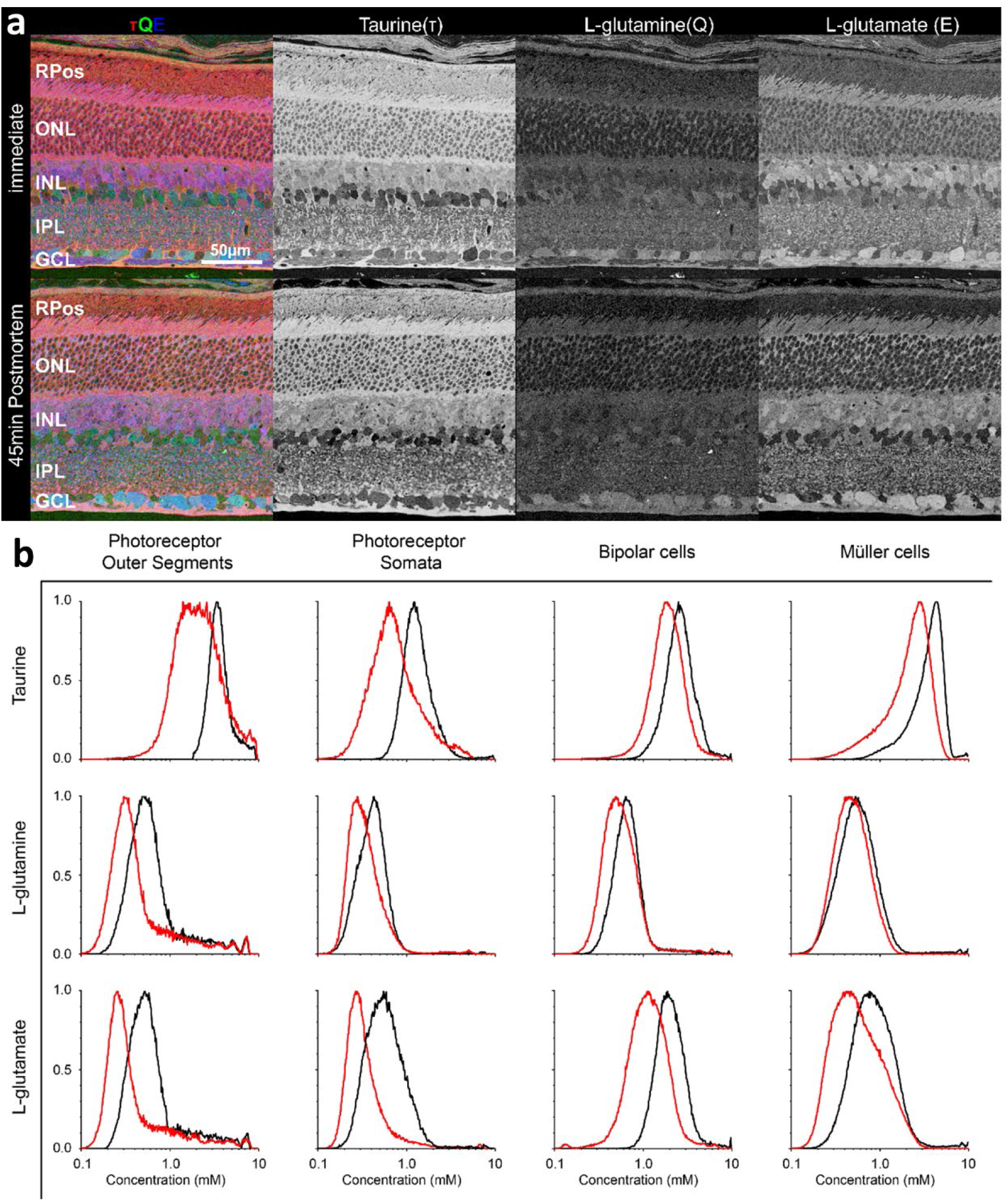
Example RGB maps of CMP staining of metabolites in the immediately fixed vs 45-minute postmortem wildtype retina **(a)**. τQE → RGB for both immediate and 45 min PM retinas. The density maps for each metabolite are also shown (greyscale). Histograms of the mean levels of Taurine, Glutamine and Glutamate measured in retinas either after immediate enucleation (black) or 45min enucleation delay (red), according to masked cell populations **(b)**.

**Extended data Figure 4.**
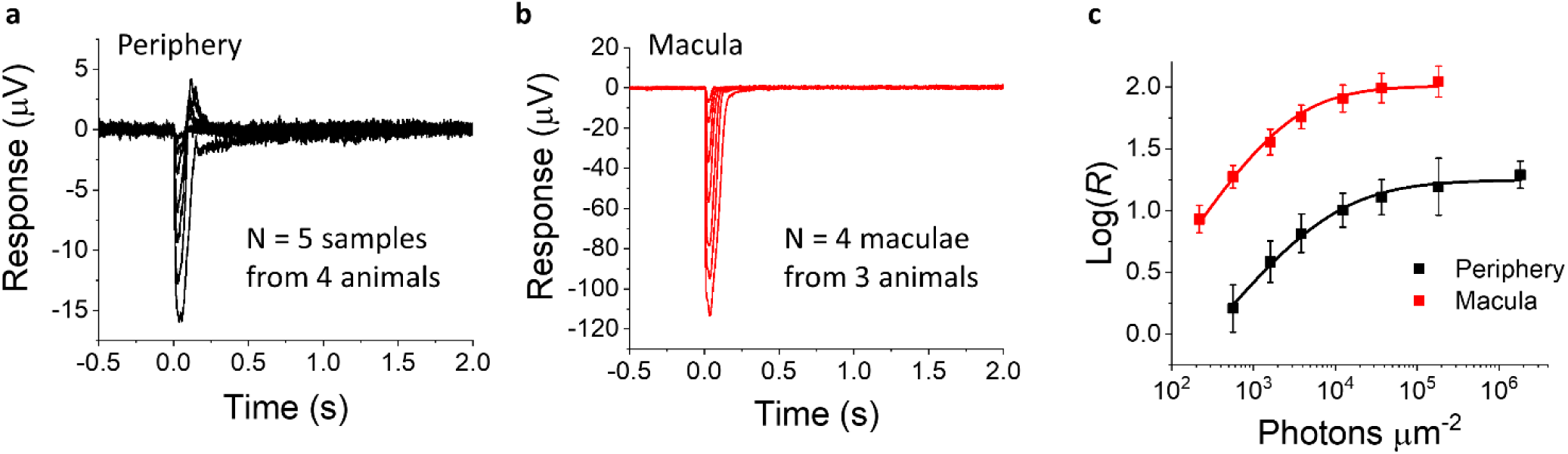
Example responses from peripheral **(a)** and macula **(b)** punches from non-human primate retinas to different light intensities from 215 to 1.8*10^6^ photons µm^-2^. Log responses plotted against light intensities used from NHP maculae (red) and peripheral retina punches (black) with Hill function (Eq. 3 in Methods) fits with I_1/2_ = 2,600 and 8,200 photons μm^-2^, respectively **(c)**. Error bars indicate ±SEM. Numbers of samples and animals are listed on plots a and b.

**Extended data Figure 5.**
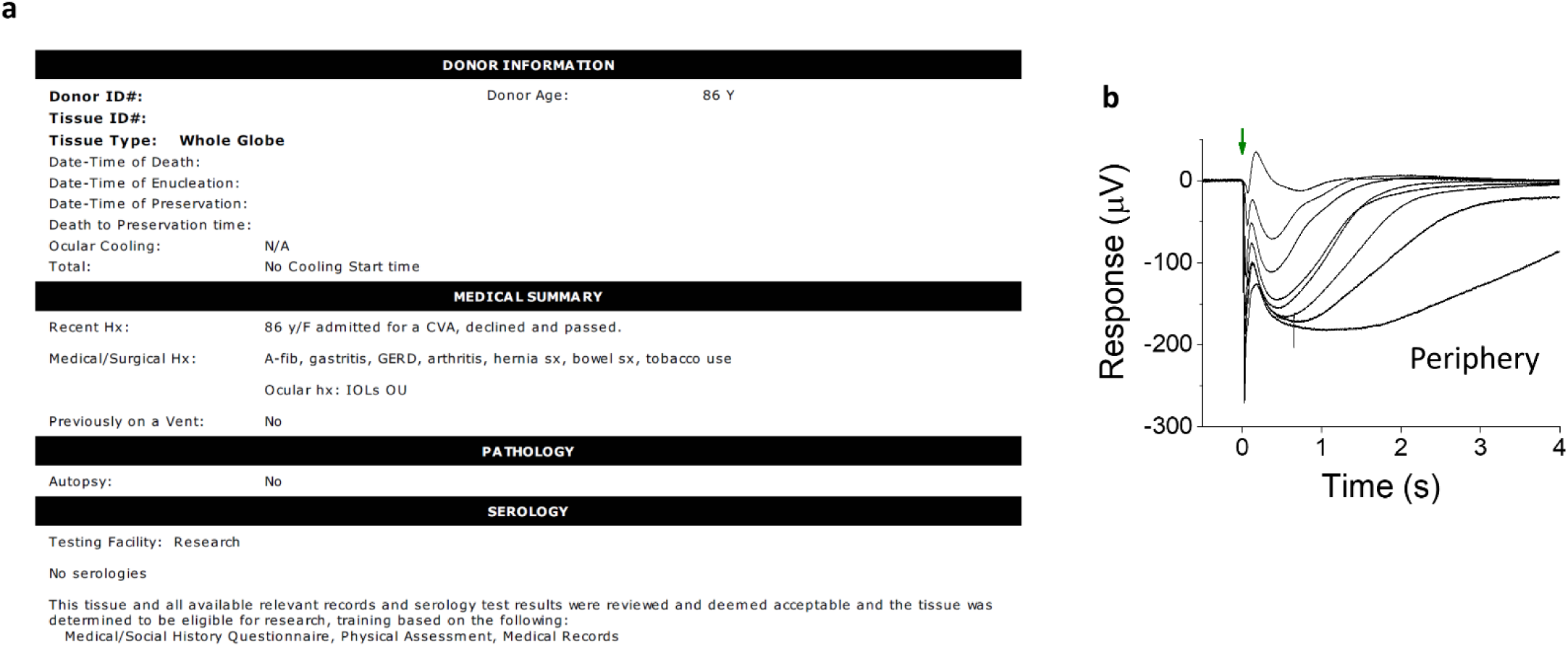
Donor information **(a)** for a single donor (donor #8) in which *ex-vivo* ERG ON-bipolar cell responses were recorded to flash intensities of 29-32,500 photons µm^-2^**(b)**. Green arrow indicates onset of flash stimuli.

